# Detection of prostate cancer in 3D pathology datasets via generative immunolabeling

**DOI:** 10.1101/2025.08.11.669726

**Authors:** Robert B. Serafin, Jennifer Salguero Lopez, Suet Chow, Rui Wang, Yujie Zhao, Elena Baraznenok, Lydia Lan, Kevin Bishop, Michelle Downes, Xavier Farre, Lawrence D. True, Priti Lal, Anant Madabhushi, Jonathan T. C. Liu

## Abstract

Recent advancements in nondestructive 3D pathology offer a complement to standard histology by enabling comprehensive volumetric analyses of intact clinical specimens (e.g. biopsies). Prior studies have demonstrated the added prognostic value of 3D pathology for prostate cancer risk stratification by correlating 3D microarchitectural features with long-term patient outcomes. However, these analyses relied on coarse manual annotations of cancer-enriched regions for downstream analysis without fine-grained delineation between often-intermixed cancerous and benign glands. To address these limitations, we have developed a 3D computational pipeline: Synthetic Immunolabeling for Generative Heatmaps of Tumor (SIGHT). SIGHT relies on deep learning-based 3D image translation models, trained in a fully supervised fashion, to convert H&E-analog 3D pathology datasets into multiplexed 3D immunofluorescence datasets that facilitate tumor detection. Our implementation of SIGHT synthetically labels two cytokeratin markers that are differentially expressed in cancerous and benign prostate glands, which are used to generate explainable 3D heatmaps of cancer-enriched regions in prostate tissues. Validation of SIGHT against ground-truth annotations from a panel of genitourinary pathologists yields an average F1 score of 0.88 which is comparable to the average inter-pathologist agreement F1 score of 0.90. To demonstrate the value of SIGHT, we developed machine classifiers of recurrence risk based on 3D glandular histomorphometric features from 75 patients. Volumetric glandular analysis in SIGHT-identified cancer-enriched regions vs. all tissue regions yields an average Kaplan-Meier hazard ratio of 3.57 (1.6 – 7.9 CI) vs. 0.92 (0.45 – 1.89 CI).

## Introduction

The management of various forms of cancer has evolved over many decades; however, the histological assessment of biopsies has largely remained as the diagnostic gold standard. In standard practice, formalin-fixed paraffin embedded (FFPE) biopsies are processed into a few thin (~5 µm thick) tissue section that are mounted onto glass slides and stained with hematoxylin and eosin (H&E) to highlight cellular nuclei and all other tissue structures. The visual interpretation of these H&E slides plays an outsized role in patient management. For example, to determine the aggressiveness of prostate cancer, PCa, which is the 2^nd^ most common cancer for men^1^, pathologists rely upon the Gleason grading system^2–4^. Gleason grading is the strongest predictor of PCa outcomes and is thus heavily relied upon for treatment recommendations. However, inter-observer variability of Gleason grading between pathologists is high^5–7^. This is in part due to Gleason grading relying upon the subjective assessment of complex branching glandular morphologies, which can be ambiguous and misleading when viewing 2D histology sections. Furthermore, pathologists review a limited number of histology sections per biopsy, typically representing less than 1% of the biopsy volume^8^. Given the dire health and financial impacts associated with the inaccurate diagnosis and grading of all cancers, including PCa, the ability to more accurately stratify patients into risk categories is of great interest.

Recent advancements in tissue-clearing techniques, high-throughput 3D microscopy, and computational tools have enabled the growing field of nondestructive 3D pathology^9–15^. Unlike standard histology, these innovations allow large clinical specimens, such as surgical excisions or biopsies, to be rapidly labeled with small-molecule fluorescent analogs of H&E and to be imaged intact without physical sectioning (**Fig. 1A, Supplementary Video 1**). Nondestructive 3D pathology generates orders-of-magnitude more data than standard 2D histology, including novel volumetric insights/features that have been shown to improve diagnostic determinations^16–19^. Previous studies have demonstrated the prognostic value of nondestructive 3D pathology by analyzing the 3D morphological characteristics of prostate biopsies and correlating them with patient outcomes. In these studies, glandular and nuclear morphologies were quantified in 1-mm core needle prostate biopsies imaged with open-top light-sheet (OTLS) microscopy. Although these studies demonstrated that 3D features outperformed analogous 2D features for patient risk stratification, these early studies relied on pathologists to manually identify cancer-enriched regions within each biopsy for downstream computational analysis. These coarse manual annotations are insufficient to provide fine-grained delineation between benign and cancerous glands that are often intermixed in prostate tissues, which may reduce the accuracy of computational classifiers^20–22^. Therefore, in this study, we sought to develop a pipeline for automated delineation of benign and cancerous regions in 3D pathology datasets (**Fig. 1B and 1C**), with glandular-scale accuracy, and to investigate if this could improve downstream prognostic analysis.

**Figure 1.**
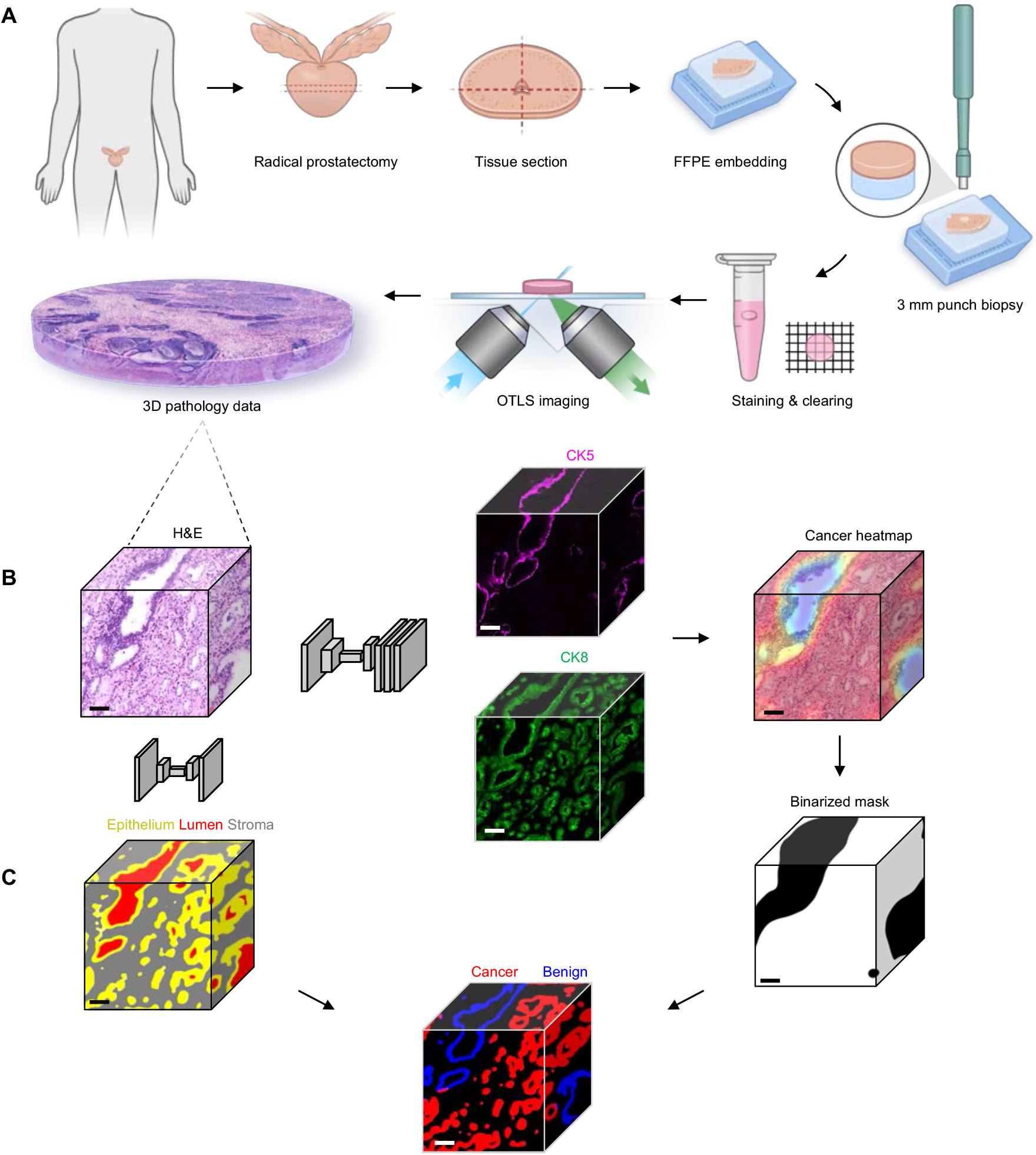
Workflow for the automated diagnosis of 3D pathology datasets using SIGHT. **(A)** 3D pathology pipeline. Radical prostatectomy specimens are formalin-fixed and paraffin-embedded (FFPE). 3-mm diameter punches are extracted, de-paraffinized, and stained with an inexpensive small-molecule (rapidly diffusing) fluorescent analog of H&E. The punches are then optically cleared and imaged using open-top light-sheet (OTLS) microscopy to produce high-resolution 3D pathology datasets. **(B)** Synthetic immunolabeling workflow. A generative adversarial network (GAN) is used to train a deep-learning model to predict 3D immunofluorescence imaging of high- and low-molecular weight cytokeratin markers (CK5 and CK8) from H&E-analog input images. This obviates the need for real immunolabeling of thick tissues, which is slow and expensive. The predicted CK5/CK8 ratio is then used to generate 3D heatmaps of cancer-enriched regions within the tissue. **(C)** Segmentation of benign vs. cancerous prostate glands. Binarized cancer heatmaps are overlaid with 3D gland segmentations (generated with a prior deep-learning model) to classify glands as benign or cancerous. Scale bars = 100 µm

In standard clinical practice, pathologists often rely on immunohistochemical staining of cytokeratin markers to differentiate between benign and malignant prostate glands^23^. Normal prostate glands consist of a bilayer of epithelial cells: a basal-cell layer, which express the high-molecular-weight cytokeratin-5 (CK5), and a luminal-cell layer, which express the low-molecular-weight cytokeratin-8 (CK8). In prostate adenocarcinoma, the most common form of PCa, the basal cell layer is lost, and cancerous glands are characterized by a single layer of luminal epithelial cells (**Fig. 2**). Although antibodies offer exceptional molecular specificity, they are expensive and diffuse very slowly in large intact tissues, such as those used for nondestructive 3D pathology. On the contrary, small-molecule fluorophores, such as the H&E-analog stains used in this study, are both cheap and rapidly diffusing in thick tissues. Synthetic immunolabeling, which uses generative deep-learning models to predict the expression of molecular targets, can bypass the need for antibody staining for both 2D and 3D pathology applications^17,24–27^. Furthermore, such “generative immunolabeling” strategies have been shown to be highly accurate for tissue structures that can be discerned by domain experts based on H&E-like images^17,27–29^. For example, we previously developed a synthetic immunolabeling pipeline, trained with a generative adversarial network (GAN), to predict the location of CK8 within 3D pathology datasets of core needle biopsies labeled with small molecule fluorescent analogs of H&E^17,30^. This, in turn, enabled automated 3D segmentation of prostate glands, all of which contain luminal epithelial cells that express CK8. We later hypothesized that this pipeline could be used to predict the expression of both CK8 (luminal cells) and CK5 (basal cells) in 3D pathology datasets of prostate biopsies, facilitating the delineation of cancerous and benign glands in an explainable way that appeals to pathologists (i.e. based on immunolabeling of standard-of-care cytokeratin biomarkers (**Fig. 1B, Supplementary Video 2**). We recognized that the development of such a computational framework, Synthetic Immunolabeling for Generative Heatmaps of Tumor (SIGHT), would enable a variety of 3D computational tasks to be fully automated for the first time (**Fig. 1C, Supplementary Videos 3 & 4**). In addition, we speculated that SIGHT could potentially improve the accuracy of these downstream analytical tasks by providing a fine-grained 3D cancer heatmap that would be impractical for human pathologists to generate at large-tissue scales.

**Figure 2.**
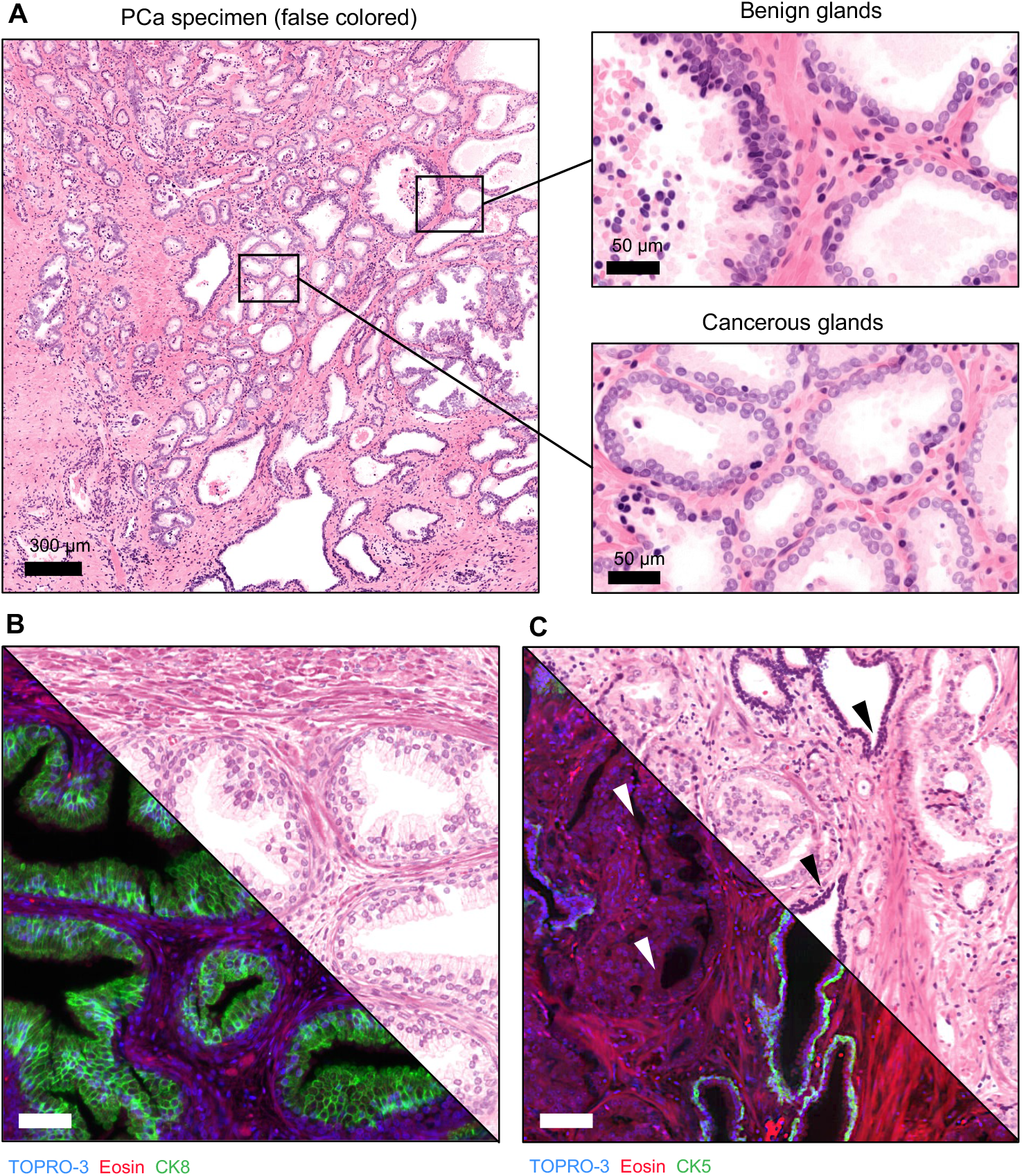
Morphological & molecular hallmarks of benign and cancerous prostate glands. **(A)** A cross-sectional view of an OTLS dataset of a 3-mm diameter prostate punch. Benign and cancerous prostate glands are often highly intermixed. Benign glands are characterized by a bilayer of epithelial cells, with an outer basal cell layer and inner luminal cell layer. Cancerous prostate glands lack a basal cell layer and are instead composed of a single layer of luminal cells. **(B)** The image on the left is an example of an OTLS dataset of benign prostate glands tri-labeled with TO-PRO-3, Eosin, and CK8 (green). CK8 is a low molecular weight cytokeratin expressed in the luminal epithelial cells that are present in all prostate glands. The virtual H&E image on the right is created by computationally false coloring the fluorescence signals from TO-PRO-3 (nuclear stain) and Eosin. **(C)** An example of an OTLS dataset of intermixed benign and cancerous prostate glands tri-labeled with TO-PRO-3, Eosin, and CK5 (green). CK5 is a high molecular weight cytokeratin expressed in the basal epithelial cells of benign prostate glands (black arrows) but is absent in the cancerous glands (white arrows).

To demonstrate the value of SIGHT-generated 3D PCa heatmaps, we analyzed 3-mm punch biopsies obtained from 75 radical prostatectomy specimens with known biochemical recurrence (BCR) outcomes. Our hypothesis was that the 3D analysis of cancer-enriched regions (automatically identified with SIGHT) vs. all tissue regions would result in significantly improved classifier performance.

## Results

### Model training and validation

To train the synthetic immunolabeling models used for automated delineation of PCa-enriched regions within 3D pathology datasets (H&E analog), we collected FFPE blocks from 15 radical prostatectomy (RP) specimens archived in an IRB-approved genitourinary biorepository at the University of Washington. Each FFPE block was sectioned into 100-µm thick slices, deparaffinized, and tri-labeled with a fluorescent analog of H&E (TOPRO-3 and Eosin) along with one of two cytokeratin antibodies (targeting CK5 or CK8). The tri-labeled slices were imaged using the high-resolution mode (~0.6 µm lateral resolution) of a 4th-generation multi-resolution OTLS system (**Fig. 3A**) ^9,15^. The tri-labeled data was used to train deep-learning models to learn the spatial relationship between H&E-analog images and cytokeratin targets. The CK5 and CK8 models were trained using a conditional generative adversarial network (cGAN)-based strategy, with 2,084 and 2,136 training ROIs, respectively (**Fig. 3B**). To validate the model performance, 35 ROIs from held-out tri-labeled specimens were processed and compared against the ground truth antibody labels. The models produced accurate synthetic immunofluorescent images, achieving average Dice coefficients of 0.72 for CK5 and 0.83 for CK8 (**Fig. 3C**). Additional details on the synthetic immunolabeling pipeline are provided in **Supplementary Fig. 2**.

**Figure 3.**
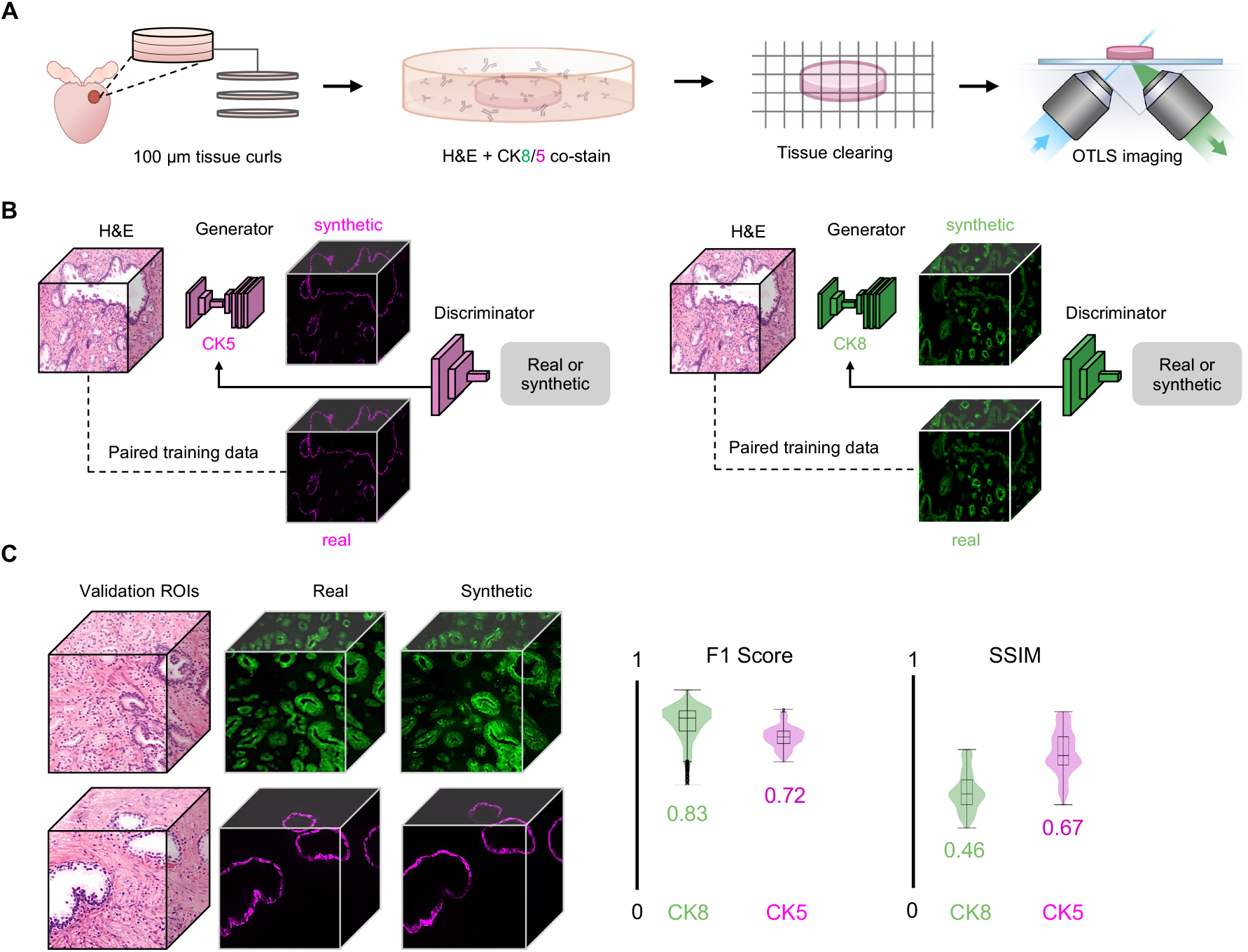
SIGHT training workflow. **(A)** For training synthetic immunolabeling models, 100-µm thick tissue sections are cut from radical prostatectomy specimens (FFPE), de-paraffinized, and tri-labeled with a fluorescent analog of H&E plus a fluorescent antibody targeting either CK5 or CK8. The specimens are then optically cleared and imaged with OTLS microscopy. **(B)** Individual models (vid2vid) are trained using conditional Generative Adversarial Networks (cGANs) to synthetically label high molecular weight cytokeratin markers (CK5) or low molecular weight cytokeratin markers (CK8) based on H&E-analog inputs. **(C)** To evaluate model performance, ROIs were held out from the training datasets and used for model validation. Each vid2vid model produced accurate synthetic images when compared to ground truth images as shown by the F1-score (dice coefficient) and the structural similarity index metric (SSIM).

### Heatmap Generation

To convert synthetic immunolabeled 3D datasets into 3D heatmaps of cancer-enriched regions, each dataset was divided into blocks corresponding to physical dimensions of 50 × 50 × 50 µm^3^. For each block, a normalized cancer-enrichment score was calculated by taking the ratio of the average intensity of CK5 to CK8 and subtracting the result from one (**Fig. 4A, Supplementary Fig. 3**). To improve the smoothness of the heatmap, the values were linearly interpolated between adjacent blocks and further refined using a Gaussian filter (**Supplementary Methods**). To validate the cancer heatmaps, we false-colored 25 specimens to mimic the appearance of standard H&E histology using a previously published algorithm^31^. From these specimens, 40 ROIs (1000 × 1000 µm in size) were randomly selected for annotation by pathologists. These regions were reviewed by a panel of three genitourinary pathologists who annotated the tumor regions using QuPath^32^ software (**Fig. 4B**). A binary mask of each pathologist’s annotations was created for each ROI. In addition, a binarized version of the SIGHT-generated heatmap was created based on a tunable threshold value applied to the cancer heatmaps. The average Dice coefficient plotted as a function of heatmap threshold value is shown in **Fig. 4C**, showing that the optimal Dice coefficient is achieved at a threshold value of 0.7. The accuracy of the automated detection algorithm (SIGHT-generated cancer heatmap) was assessed by calculating the recall, precision, and F1 score (Dice coefficient) of the SIGHT-generated heatmap vs. the pathologists’ annotations (**Fig. 4D**). The average F1 score between the heatmaps and pathologist annotations is 0.88, which is comparable to the average inter-pathologist F1 score of 0.90 (**Fig. 4E**). In other words, the accuracy of our 3D cancer heatmaps approaches the agreement between any two experienced genitourinary-specialist pathologists.

**Figure 4.**
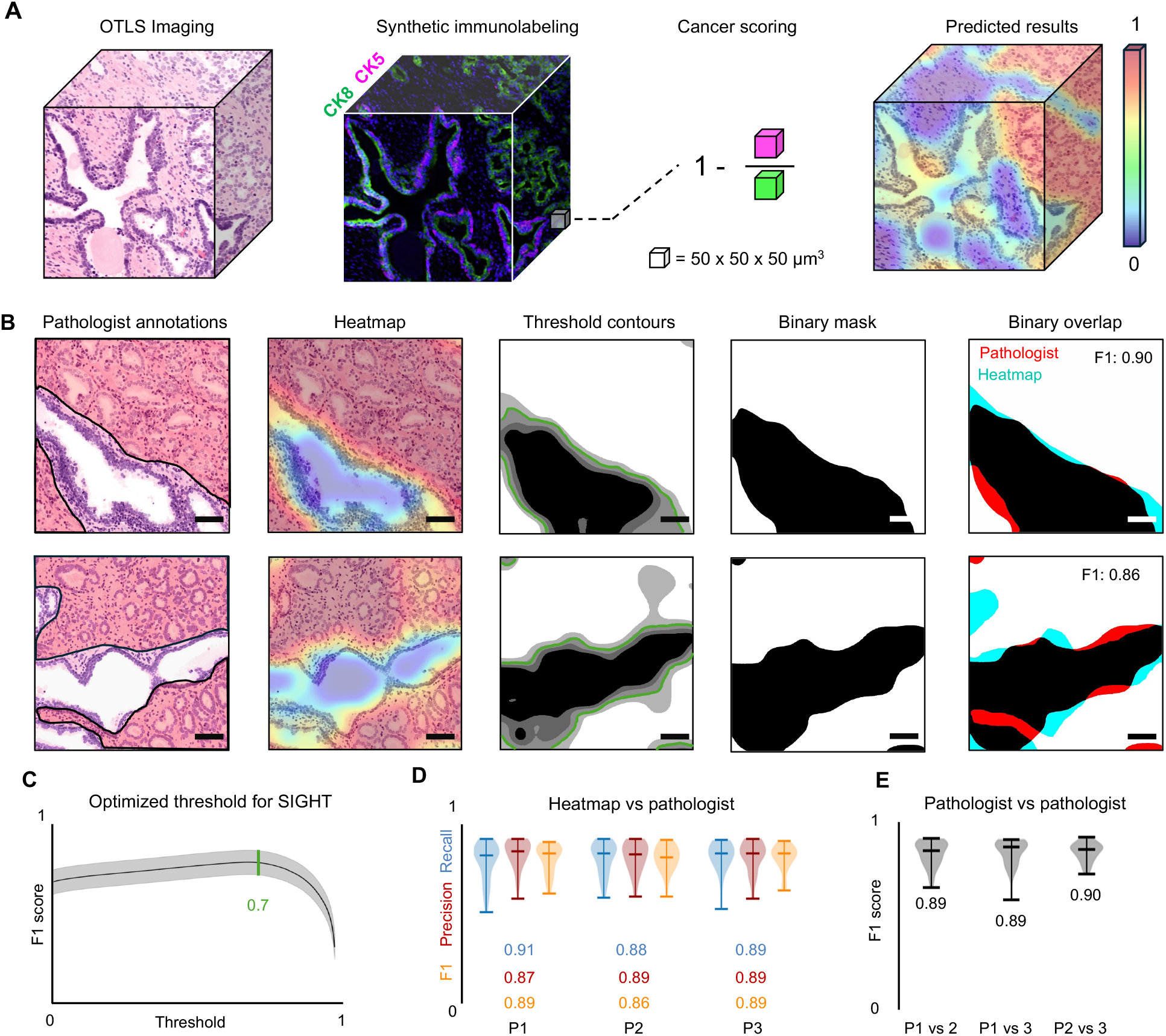
SIGHT workflow and validation. **(A)** Overview of the SIGHT workflow for automated cancer detection in 3D pathology datasets. Synthetic CK5 and CK8 datasets are used to generate cancer heatmaps for small sub-volumes measuring 50 × 50 × 50 µm in size, where the “cancer-enrichment” score is the ratio of CK5 to CK8 expression (averaged within each sub-volume) subtracted from one. **(B)** To validate SIGHT predictions, three genitourinary pathologists independently annotated cancerous glands contained in 40 regions of interest. **(C)** A range of threshold values were applied to the corresponding SIGHT heatmaps to create a binary cancer segmentation mask, which was compared to the expert annotations to identify the optimal threshold where the F1 score (Dice coefficient) was maximized. **(D)** Once the optimal threshold was established (0.7), binarized synthetic (SIGHT) heatmaps were evaluated against the pathologist annotations through several accuracy metrics: recall, precision, and F1. SIGHT achieves an average recall of 0.90, precision of 0.89, and F1 score of 0.88. **(E)** The average inter-pathologist agreement is calculated as F1 = 0.89, showing that SIGHT performs at a comparable level to expert human observers. Scale bars = 100 µm

To evaluate whether SIGHTs segmentation of cancer-enriched regions was statistically equivalent to that of expert pathologists, we applied the two one-sided test (TOST) procedure using paired F1 scores. Equivalence bounds were set to ±1 standard deviation of the F1 scores computed across all pairwise pathologist annotations, capturing the inter-pathologist variability. The TOST assessed whether the mean difference in F1 scores between the SIGHT heatmaps and each pathologist entirely within these bounds p = 0.012, 0.0091, and 0.017 respectively, supporting statistical equivalence to expert-level performance.

### Imaging and analysis workflow for clinical-validation cohort

To demonstrate the value of the 3D cancer heatmaps (SIGHT) for improving the accuracy of machine classifiers of recurrence risk, a clinical-validation study was performed using archived radical prostatectomy (RP) specimens. Our study consisted of N=75 patients with known time-to-recurrence outcomes. Most of the patients selected for this cohort belonged to Gleason grade groups 1 to 3, for which there is often difficulty in determining treatment strategy (e.g. active surveillance vs. curative surgery or radiation). Additional details on the patient cohort are show in **Supplementary Table 1**. One 3-mm diameter punch was extracted from the index lesion for each patient. Following deparaffinization of the FFPE tissue specimens, the 3-mm punch were stained with an inexpensive and rapidly diffusing small-molecule fluorescent analog of H&E, optically cleared with a simple dehydration and solvent-based clearing protocol to render them transparent to light, and then imaged with a high resolution open-top light-sheet (OTLS) microscopy platform to obtain whole-specimen 3D pathology datasets^9^ (**Fig. 1A**).

Each 3D pathology dataset was processed using SIGHT, with the H&E-analog fluorescence channels serving as model inputs (**Supplementary Fig. 2, Supplementary Video 2**). The high molecular-weight cytokeratin CK5 is expressed in the basal cell layer of prostate glands, which are absent in cancer glands. The low molecular-weight cytokeratin CK8 is expressed in the luminal cell layer present in all prostate glands. The synthetic cytokeratin images were used to generate a 3D heatmap of the cancer-enriched regions of each 3-mm diameter tissue punch (**Fig. 1B and 4A, Supplementary Fig. 3, Supplementary Video 3**). 3D segmentation masks of the glandular epithelium, lumen, and stromal regions of each punch were generated using a previously published deep-learning based 3D segmentation pipeline (**Supplementary Fig. 4**)^33^. Cancer-enriched and benign regions of each dataset were segmented by thresholding the heatmaps at an optimal value (**Fig. 1C and 4C**). Combining the 3D gland segmentations with the binary segmentation mask of the cancer-enriched regions allowed us to classify glandular epithelia as either cancerous or benign (**Supplementary Video 5**).

3D histomorphometric features of the glandular epithelium, lumen, and stroma were quantified across the whole biopsy, across cancer-enriched regions alone, or across benign region alone. A full list of histomorphometric features is provided in **Supplementary Table 2**. For each condition (i.e. whole biopsy or cancer-enriched or benign), we implemented a 10-fold cross-validated survival analysis using a Cox regression model based on 5-year time-to-biochemical recurrence (BCR) outcomes. For each fold, we calculated risk scores and divided patients into two risk groups. The cutoff point between high and low risk was set using the median score from the training data and then used to classify patients in the test data. KM-curves of BCR-free survival indicate that models trained on 3D features extracted from the whole punch biopsy (mixture of cancer and benign glands) failed to stratify patients into high- and low-risk groups: p > 0.05, C-index = 0.522, and HR = 0.92 (**Fig. 5**). Models trained on 3D features from SIGHT-identified cancer-enriched regions achieved statistically significant risk stratification (p < 0.05, C-index = 0.653, and HR = 3.57). Classification models based on benign regions were not prognostic.

**Figure 5.**
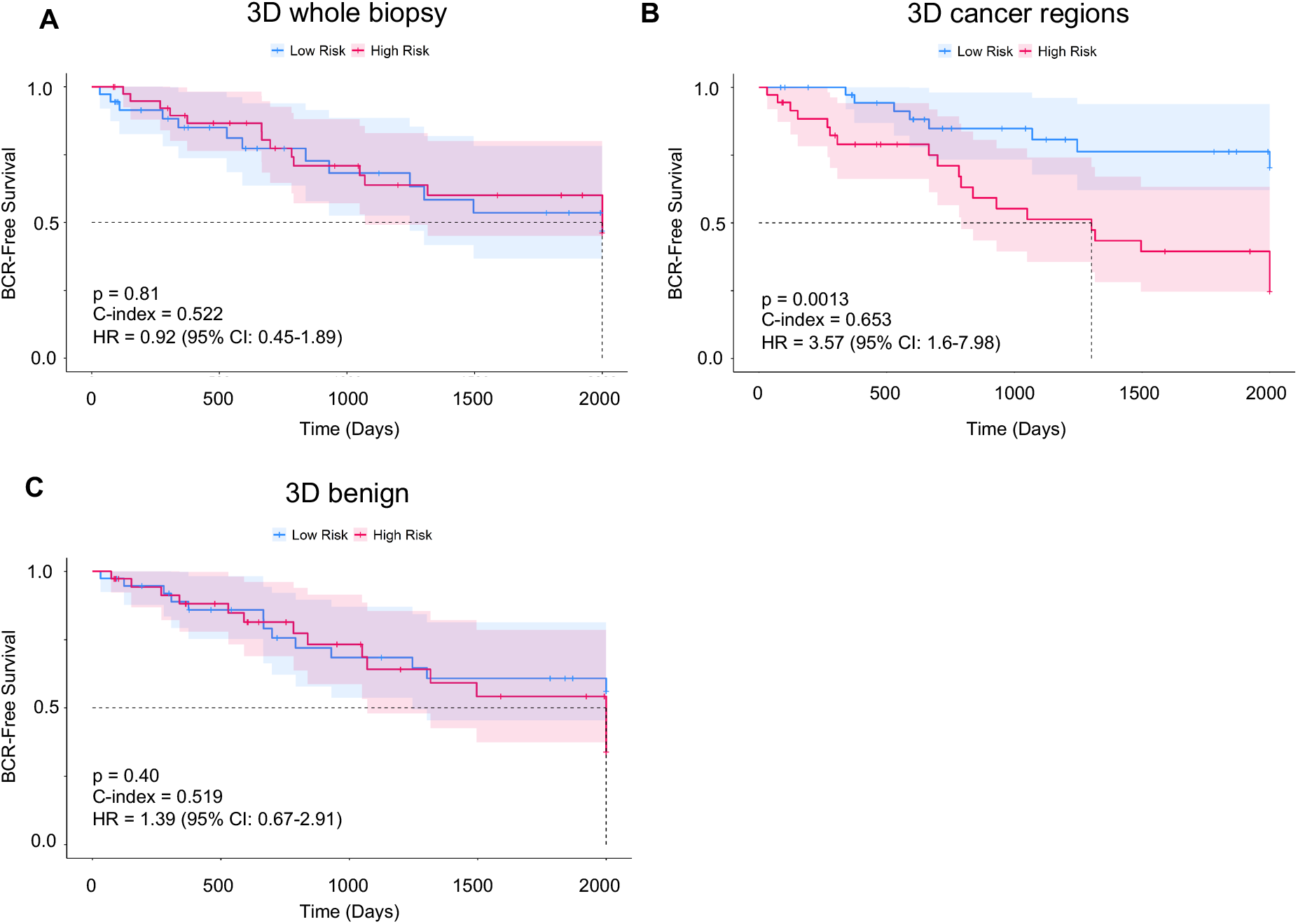
Risk stratification based on SIGHT-identified cancer regions in 3D pathology datasets. Using 5-year time-to-biochemical recurrence (BCR) outcomes for 75 patients, Cox regression models with 10-fold cross-validation were used to evaluate the prognostic value of features extracted from **(A)** whole biopsy volumes, **(B)** SIGHT-identified cancer-enriched regions or **(C)** benign regions. Features derived from SIGHT-predicted cancer regions were prognostic for BCR-free survival, whereas features extracted from the entire biopsy volume or from benign regions were not prognostic.

## Discussion

In recent years, there has been a push to translate 3D high-resolution imaging modalities beyond traditional research applications and into clinical settings. However, considering the massive size and complexity of such high-resolution 3D pathology datasets, there is an increasing need for computational tools to assist clinicians with their interpretation and to facilitate clinical decision-making. Amongst various computational approaches, there is a recognized value in incorporating intuitive and explainable tissue features, grounded in existing clinical knowledge, for the analysis of specific use cases. For example, recent studies have demonstrated the benefits of computational analyses of 3D glandular architectures and nuclear morphologies for prostate cancer prognostication^16–18^. However, these early studies relied on coarse manual annotations by pathologists to delineate cancer-enriched regions for downstream 3D analysis. This manual approach was inefficient and subjective and could not provide fine-grained delineation of cancer-enriched vs. benign regions, which are often intermixed in human specimens. Given these limitations, we have developed Synthetic Immunolabeling for Generative Heatmap-based Tumor detection (SIGHT), an automated 3D cancer detection platform. SIGHT produces a 3D heatmap of cancer enrichment within 3D pathology datasets of tissues labeled with and inexpensive and rapidly diffusing fluorescent analog of H&E. Importantly, SIGHT bypasses the need for tedious and subjective manual 3D annotations by pathologists and is trained with objective molecular biomarkers of prostate cancer. Furthermore, SIGHT leverages the speed and efficiency of deep learning to overcome the practical limitations of immunolabeling thick tissues with expensive and slowly diffusing (large) antibodies.

In the context of identifying malignant (cancerous) prostate glands, SIGHT uses deep learning-based 3D image translation, trained via a generative adversarial network (GAN) paradigm, to synthesize images of high- and low-molecular weight cytokeratin markers (CK5 and CK8) that are differentially expressed in benign and cancerous prostate glands. Each image-translation model (for CK5 or CK8) was trained on 3D OTLS microscopy datasets of tri-labeled prostate tissues, in which tissues were co-stained with our fluorescent analog of H&E (model inputs) plus a targeted antibody of interest (either CK5 or CK8 in this case) as the desired model output. By calculating the ratio of CK5 to CK8 expression, SIGHT generates 3D heatmaps of cancer-enriched regions within intact prostate tissues with glandular-level granularity. Notably, when benchmarked against ground-truth annotations by expert genitourinary pathologists, SIGHT’s predictions of cancer regions showed agreement levels that approach the interobserver agreement amongst the pathologists themselves. This underscores SIGHT’s potential to achieve expert-level performance in prostate cancer delineation, but in a fully automated way that can scale to massive 3D pathology datasets. To evaluate the potential of SIGHT for improving prostate cancer risk stratification, we applied SIGHT to OTLS datasets from 75 patients with known time-to-event outcomes. Our findings suggest that survival models trained on features extracted from SIGHT-identified cancer regions are better at predicting patient outcomes than models trained on features extracted from the whole biopsy.

There are a limited number of generative network architectures designed for 3D image translation. SIGHT currently relies on GANs that are advantageous for their fast inference speed in high-volume 3D pathology applications. However, GANs can be difficult to train due the requirement for large and diverse datasets. Diffusion models could serve as a viable alternative for achieving higher-fidelity synthetic immunolabeling to enhance future iterations of SIGHT^39^. Note that once sufficient synthetic immunolabeled datasets are generated and validated, a direct segmentation model (i.e. H&E-analog to cancer heatmap) could be trained^33^, which would be faster and potentially more generalizable. In other words, SIGHT is an objective (chemically based) annotation-free segmentation method that could enable the training of more-efficient direct segmentation models in the future. Nevertheless, the ability to generate synthetic immunolabeling datasets, which replicate the appearance of standard clinical IHC stains that pathologists already trust and rely upon in their daily practices, is an attractive aspect of SIGHT that could facilitate clinical adoption.

There are several opportunities to leverage the initial success of SIGHT. First, in this work we primarily focused on prostate cancers exhibiting Gleason patterns 3 and 4, for which a major clinical challenge is to determine which patients would benefit from curative therapies such as surgery or radiation. SIGHT’s capabilities could be expanded to include other morphologies (e.g. pattern 5 glands) during training. Second, training additional models to predict other cancer biomarkers could enhance the sensitivity and specificity of SIGHT for delineating cancer vs. benign regions ^40,41^, or to identify cancer subtypes that could be targeted with pharmacological treatments^42,43^. The combination of additional molecular targets and morphological primitives (e.g. vessels, nerves, collagen, immune cells), identified with SIGHT, could enable the development of powerful risk classifiers and predictive assays for a variety of diseases.

Outside of prostate cancer, SIGHT could be adapted to other use cases where there is a known association between high-resolution histomorphology and molecular expression, For example, recent studies have attempted to predict HER2 expression in breast cancer specimens without immunolabeling based on label-free autofluorescence or phase-contrast microscopy^27–29,44^, modalities that primarily capture tissue morphology. Similarly, SIGHT could integrate with other molecular profiling techniques beyond immunolabeling, including spatial transcriptomics, multiplexed protein assays, or other molecular signatures that distinguish different cell and tissue types with high specificity.

In summary, SIGHT is a powerful paradigm for the segmentation of various histologic primitives, including cancer-enriched regions, in an objective and annotation-free manner. We show that this is especially valuable for applications of 3D pathology in which manual annotations are prohibitive and where immunolabeling is slow and costly. Our current implementation of SIGHT, reported here for the automated 3D delineation of prostate cancer-enriched tissue regions, is a critical step towards establishing, for the first time, a fully automated computational pipeline for risk stratification based on the analysis of interpretable hand-crafted 3D features. This fully automated pipeline will enable scaled-up studies on hundreds to thousands of patient specimens to convincingly demonstrate the value of 3D vs. conventional 2D histopathology for guiding critical treatment decisions for cancer patients.

## Materials and methods

### Tissue preparation

Tissue punches (3-mm diameter) were obtained from FFPE blocks of radical prostatectomy (RP) specimens and deparaffinized in xylene (Fisher Scientific, cat. no. X3P-1GAL) for 48 hours at 60°C. Following deparaffinization, the punches were washed twice in 100% ethanol (Decon Laboratories, cat. no. 2701) for 1 hour^9^. For H&E-analog staining, nuclei were stained with TOPRO-3 Iodide (Cat: T3605, Thermo-Fisher) at a 1:500 dilution and Alcoholic Eosin Y 515 (Cat: 3801615, Leica Biosystems) at a 1:100 dilution for 48 hours with light agitation. Tissues were stained in 1.5 mL of a 70% ethanol, 30% DI water buffer with 10 mM NaCl (Spectrum Chemical, cat. no. SO160) adjusted to pH 4 (+/-0.05) by adding HCL (Fisher Scientific, cat. no. SA54-1). After staining, tissues were dehydrated in 100% ethanol for two 1-hour increments. Tissues were cleared in ethyl cinnamate (Cat: AAA1290622. Fisher Scientific) at room temperature overnight.

### Tri-labeling

A detailed overview of the tri-labeling procedure is provided in **Supplementary Fig. 1**. 100-micron thick prostate curls were cut from 15 FFPE blocks of RP specimens from a cohort of patients originally diagnosed with low-to intermediate-risk PCa, i.e. Gleason grade groups 1 (3 + 3), 2 (3+4) and 3 (4+3). Tissue curls were deparaffinized over 48 hours in 60°C xylene (Fisher Scientific, cat. no. X3P-1GAL), followed by rehydration through an ethanol-water gradient: 100%, 75%, 50%, and 25% ethanol (Decon Laboratories, cat. no. 2701) and a final transfer to PBS for 1 hour. Samples were then incubated in Alexa Fluor 488 NHS Ester (Invitrogen/Thermo Fisher Scientific A20000) at 5 µg/mL overnight at 37°C. Subsequent washing, permeabilization, and blocking were completed using PBS/0.2% Triton X-100, 20% DMSO, 0.3M glycine, and 6% donkey serum for three days. Following the blocking step, samples were incubated in mouse anti-human CK5 primary antibody (Thermo Fisher Scientific MA5-12596, Abcam ab17130) or mouse anti-human CK8 primary antibody (Thermo Fisher Scientific MA5-14088, Abcam ab17139) at 1:20 dilution in a PBS solution with 0.2% Tween-20, 10 µg/ml heparin, 3 mM NaN3, 5% DMSO and 3% donkey serum at 37°C, followed by washing and incubation with Alexa 647 donkey anti-mouse secondary antibody (Jackson ImmunoResearch Laboratories 715-605-150) at a 1:100 dilution. Nuclear staining was achieved using SYTO 85 (5 µM) overnight at 37°C. The samples were dehydrated through a reverse ethanol gradient and cleared in ethyl cinnamate (Cat: AAA1290622. Fisher Scientific) for two 30-minute intervals.

### Imaging and post processing

H&E analog-stained 3-mm prostate punches were imaged on an OTLS microscope. Samples were illuminated using 561 nm (eosin) and 638 nm (TO-PRO3) excitation wavelengths (Skyra, Cobolt Lasers, Hubner Photonics, Kasel, Germany). For tri-labeled specimens (100-micron thick), imaging was performed using 488 nm (NHS Ester), 561 nm (SYTO 85), and 638 nm (CK5 or CK8) excitation wavelengths. Continuous (stitched and fused) 3D volumes were created using the BigStitcher^35^ plugin for ImageJ with 2X down sampling along each dimension such that pixel spacing for the whole volume was 0.572 µm/pixel. 3D volumes were flat fielded (stripe correction and depth adjustment) and intensity normalized using a previously published post-processing method^9^.

### SIGHT model training

We extracted training ROIs from the fused tri-labeled datasets. Training ROIs were 512 × 512 × 50 pixels in size at a sampling pitch of 0.572 µm/pixel. The tri-labeled data was separated into individual channels for training, such that *vid2vid*^30^ could learn to predict the expression of each molecular target based on the corresponding H&E-analog images. The CK5 and CK8 models were trained with 2,084 and 2,136 training ROIs, respectively. See **Supplementary Information** for additional details.

### SIGHT data preparation, inference, and volume assembly

3D pathology datasets of 3-mm diameter punches are ~ 90 GB in size after standard post-acquisition processing^9^. To accommodate for the size of each dataset, the procedure for running the SIGHT pipeline is outlined in **Supplementary Fig. 2**. First, the H&E analog channels are broken into 25 separate blocks (1024 × 1024 × 640 voxels in size with 25% overlap). Each image block is written to disk as a z-stack (separate individual jpegs per z plane) using the directory architecture expected by *vid2vid*. The H-analog channel (TO-PRO3) is used as two input channels ch0 & ch1, while the Eosin channel is used as the third input channel, ch2. Following inference by *vid2vid*, the synthetic image blocks are reassembled into a single 3D image volume using the stitching plugin from ImageJ^36^

### Heatmap generation

To detect the cancer-enriched regions of 3D pathology datasets, SIGHT follows the sequence of steps outlined in **Fig. 4A** and **Supplementary Fig. 3**. Briefly, OTLS datasets are broken up into individual cubes measuring approximately 50 µm per side. A ‘cancer score’ is calculated for each cube by calculating the ratio of the average CK5 signal (above background) to the average CK8 signal (above background) within each 50-µm cube, and subtracting if from one:

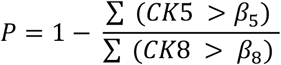

Here, CK5 and CK8 represent the average signal levels of each synthetic immunolabel within the volume, and β_5_ and β_8_ denote their respective background thresholds.

Following the initial scoring procedure, the heatmap is relatively coarse. To smooth the heatmap, a median filter with a cross-shaped structuring element (connectivity = 1) is applied across each block. Next, scores between adjacent blocks are smoothed through linear interpolation followed by a gaussian filter (σ = 25 voxels). The scoring procedure is implemented using the Python libraries *numpy* and *scikit-image*. For efficient processing, the framework leverages the automatic parallelization capabilities of the *dask* library.

### Comparing heatmap segmentations with pathologist annotations

25 specimens with a high degree of intermixing between benign and cancerous glands were chosen for annotation by three expert GU pathologists. To aid pathologist interpretation and annotation, H&E analog channels from each dataset were false colored to mimic the standard appearance of H&E histology^31^. 40 regions of interest, each measuring 1 mm × 1 mm in size, were randomly selected for annotation. Pathologists were instructed to annotate the boundary between benign and cancerous regions of each image using Qupath (**Fig. 4B**).^32^ Annotations were converted into binary masks using the *geopandas* python library. Annotations of each ROI were compared to binarized SIGHT heatmaps by measuring the Dice coefficient (F1 score). Optimal thresholds for binarizing the heatmaps were then determined to maximize the average F1 score (i.e. using SIGHT to mimic the pathologist’s annotations as closely as possible).

### Glandular segmentation and feature extraction

3D gland segmentations of our 3D pathology datasets of prostate cancer were generated using a previously published direct deep learning model trained with nnU-Net ^33,37^. Using the 3D gland segmentations, 14 features related to gland size and shape were calculated. These features, along with brief descriptions are listed in **Supplementary Table 2**. The first feature set quantified the volumetric extent of tissue compartments, including the lumen, epithelium, and stroma. Additional feature sets described compartment shapes using triangle mesh approximations. Examples include surface-area-to-volume ratio; average 3D surface curvature; and gland-to-convex-hull ratio (G/H), defined as the volume (3D) ratio of the gland mask (epithelium + lumen) to its circumscribing convex hull. Features were extracted using the *scikit-image* and *scipy* python libraries, specifically the *regionprops, mesh_surface_area*, and *ConvexHull* methods. Measurements of glandular and luminal curvature were acheived using the discrete_gaussian_curvature_measure method from the *trimesh* library using meshes generated via the *marching_cubes* algorithm in *scikit-image*.

### Prognostic analysis

For each tissue compartment (whole biopsy, cancer-enriched region, and benign region), a 10-fold cross-validated survival analysis was performed using a Cox regression model in MATLAB. Risk scores were calculated for each fold, and patients were stratified into high- and low-risk groups. The median risk score from the training data was used as the threshold to differentiate between high- and low-risk groups, and this threshold was subsequently applied to classify patients in the test data. The model performance was quantified with p-values (by log-rank test), hazard ratios (HRs) and concordance index (C-index) metrics.

## Ethics Statement and Patient Consent

Radical prostatectomy specimens used in this study were archived in IRB-approved genitourinary biorepositories at the University of Washington and University of Pennsylvania

## Author contributions

**R. Serafin**: Conceptualization, data curation, software, formal analysis, validation, investigation, visualization, methodology, writing-original draft, writing-review and editing. **J. S. Lopez:** data curation, software, formal analysis, methodology. **S. Chow:** data curation, software, formal analysis, tissue preparation. **R. Wang:** Software, methodology. **E. Baraznenok**: data curation, tissue preparation. **L. Lan**: data curation, tissue preparation. **K. Bishop:** Data curation. **M. Downes:** data curation. **X. Farre:** data curation. **L. True:** data curation, methodology, formal analysis, validation, investigation. **P. Lal**: data curation, validation, writing-review and editing. **A. Madabhushi**: Resources, formal analysis, supervision, funding acquisition, project administration, writing-review and editing, **J.T.C. Liu:** Conceptualization, resources, formal analysis, supervision, funding acquisition, validation, investigation, visualization, writing-original draft, project administration, writing-review and editing.

## Acknowledgements

Research reported in this publication was supported by ARPA-H under contract number D24AC00357 (Liu) and D25AC00140 (Madabhushi). Research was also supported by the National Institute of Biomedical Imaging and Bioengineering (NIBIB) under R01EB031002 (Liu), the National Institute of Diabetes and Digestive and Kidney Diseases (NIDDK) under R01DK138948 (Liu), the National Cancer Institute (NCI) under R01CA268207 (Liu and Madabhushi), R01CA268287 (Madabhushi), U01CA269181(Madabhushi), R01CA249992 (Madabhushi), R01CA202752 (Madabhushi), R01CA208236 (Madabhushi), R01CA216579 (Madabhushi), R01CA220581 (Madabhushi), R01CA257612 (Madabhushi), 1U01CA239055 (Madabhushi), 1U01CA248226 (Madabhushi), 1U54CA254566 (Madabhushi), the National Heart, Lung and Blood Institute (NHLBI) under 1R01HL15127701A1 (Madabhushi) and R01HL15807101A1 (Madabhushi), VA Merit Review Award IBX004121A (Madabhushi) from the United States Department of Veterans Affairs Biomedical Laboratory Research and Development Service the Office of the Assistant Secretary of Defense for Health Affairs, and sponsored research agreements from Bristol Myers-Squibb, and Astrazeneca. Additional support was through the Department of Defense (DoD) Prostate Cancer Research Program (PCRP) grant W81XWH-18-10358 (Liu and True), W81XWH-14-2-0183 (True), W81XWH-20-1-0851 (Madabhushi and Liu), the Pacific Northwest Prostate Cancer SPORE P50CA97186 (True), and the Canary Foundation. The content is solely the responsibility of the authors and does not necessarily represent the official views of the National Institutes of Health, the U.S. Department of Veterans Affairs, the Department of Defense, or the United States Government.

