## Supplemental Materials for "Detection of prostate cancer in 3D pathology datasets via generative immunolabeling"

### Supplementary methods

Collection and processing of prostate tissues

Training synthetic immunolabeling models

SIGHT inference pipeline

SIGHT heatmap generation

3D Gland segmentation

Separation of benign and cancerous prostate glands

### Supplementary Figures

Tri-labeling protocol for OTLS datasets

Synthetic immunolabeling inference workflow

Heatmap generation from synthetic immunolabels

3D Gland segmentation workflow

**Supplementary Tables**

Relevant clinical parameters for study cases (N=75)

List of glandular features

**Supplementary Videos**

Nondestructive 3D pathology dataset of a 3 mm punch biopsy of prostate tissue

Fly through of synthetically immunolabeled prostate biopsy

3D gland segmentations following application of SIGHT heatmap

Depth stack visualization of SIGHT-generated heatmaps overlaid on OTLS microscopy data, false-colored to resemble H&E histology.

Classification of cancerous and benign prostate glands

**Collection and processing of prostate tissues**

To train the CK8 and CK5 synthetic immunolabeling models, we collected one FFPE block from each of 15 radical prostatectomy (RP) specimens archived in an IRB-approved genitourinary biorepository at the University of Washington (UW). Based on the original pathology reports generated from the RP specimens, 7 specimens were from Gleason Grade 3+3 and 8 specimens from Gleason Grade 3+4 and 4+3. The imaging data from this cohort allowed us to train an image-translation model for low- to intermediate-risk PCa (Grade Group 1-3). We used a vibratome to cut each block into 100 µm sections. This thickness provided sufficient 3D context to train and test the synthetic immunolabeling models while allowing for uniform antibody diffusion and staining within the tissue slices. FFPE sections were deparaffinized in 60°C xylene for 48 hours.

A hotplate with magnetic stirrer was used to maintain the temperature of the xylene and to promote fluid convection around the specimens.

Our tri-labeling protocol (originally adapted from the iDISCO protocol) was the same as our prior work in synthetic immunolabeling. Details are shown in **Supplementary Figure 1**. Deparaffinized tissue curls were incubated in Alexa Fluor 488 NHS Ester (Invitrogen/Thermo Fisher Scientific A20000) at 5 μg/mL overnight at 37°C. Labeling the tissue with NHS Ester (which binds to all proteins) prior to immunostaining is crucial to prevent unbiased training of the synthetic immunolabeling models as it prevents NHS Ester from binding to the CK8 or CK5 antibodies. Subsequent washing, permeabilization, and blocking were completed using PBS/0.2% Triton X-100, 20% DMSO, 0.3M glycine, and 6% donkey serum for three days. Following the blocking step, samples were incubated in mouse anti-human CK5 primary antibody (Thermo Fisher Scientific MA5-12596, Abcam ab17130) or mouse anti-human CK8 primary antibody (Thermo Fisher Scientific MA5-14088, Abcam ab17139) at 1:20 dilution in a PBS solution with 0.2% Tween-20, 10 μg/ml heparin, 3 mM NaN3, 5% DMSO and 3% donkey serum at 37°C, followed by washing and incubation with Alexa 647 donkey anti-mouse secondary antibody (Jackson ImmunoResearch Laboratories 715-605-150) at a 1:100 dilution. Nuclear were stained with SYTO 85 (5 μM) overnight at 37°C. The samples were dehydrated through a reverse ethanol gradient and cleared in ethyl cinnamate (Cat: AAA1290622. Fisher Scientific) for two 30-minute intervals.

For tri-labeled specimens, imaging was performed using 488 nm (NHS Ester), 561 nm (SYTO 85), and 638 nm (CK5 or CK8) excitation wavelengths, with 10 ms exposure. Continuous (stitched and fused) 3D volumes were created using the BigStitcher plugin for ImageJ with 2X down sampling along each dimension such that pixel spacing for the whole volume was 0.572 μm/pixel.

**Training synthetic immunolabeling models**

GANs offer a flexible approach to image translation by eliminating the need for manually designed models and loss functions, which traditionally require domain-specific expertise. A GAN consists of two independent networks: the discriminator, which aims to distinguish between real and generated (fake) images, and the generator, which aims to produce authentic-looking data capable of deceiving the discriminator. As an extension of the GAN structure, conditional GANs (cGANs) enable models to learn to map input images to outputs in a fully supervised manner by conditioning both the generator and discriminator on prior information. In this case, the tri-labeled OTLS datasets provide the required paired images, where the H&E analog channels serve as model inputs and immunofluorescence channel (CK8 or CK5) as outputs

For SIGHT, we built upon our previous work in synthetic immunolabeling. To ensure continuous and realistic 3D inference, image volumes must first be 'primed' with a 2D image-translation model on the initial levels. Following our established approach, we used an adapted version of *pix2pix*, a cGAN originally designed for image-to-label translation and widely adopted in biomedical imaging. This version of *pix2pix* uses 9-layer U-Net generator and a PatchGAN discriminator to learn a direct mapping between H&E analog inputs and their corresponding cytokeratin outputs (CK8 or CK5). For 3D image translation we used vid2vid, a video-translation model that ensures continuity between sequential frames. In this case, we treated 3D images as a "z-stack" of 2D images, allowing the model to maintain spatial consistency along the depth dimension. The model also employed a multi-scale PatchGAN discriminator for individual images and a sequence discriminator to enforce consistency across the 3D spatial domain.

Training datasets for each synthetic immunolabeling models were screened/curated primarily based on the image quality (i.e. evenness of staining and signal to noise ratio) of the CK8 or CK5 immunolabeling. The GANs used for synthetic immunolabeling (pix2pix & vid2vid, details below) were originally designed to process 8-bit RGB images, whereas our OTLS datasets were acquired using a 16-bit sCMOS camera. To ensure optimal utilization of the 8-bit dynamic range in both the training images and synthetic immunolabeling results we implemented a percentile-based histogram adaptation as a preprocessing step. To minimize loss of signal during the 8-bit conversion process, the $2^{nd}$ and $98^{th}$ percentiles of the 16-bit image data was mapped to the minimum and maximum values of the 8-bit range.

Individual pix2pix and vid2vid models were trained for each cytokeratin target. The final training dataset for the vid2vid models consisted of 2136 tri-labeled ROIS for CK8 and 2084 tri-labeled ROIs for CK5, each containing 50 levels. For pix2pix the training data consisted of 5 individual 2D levels, separated by 10 µm, taken from the 3D image sequences of the vid2vid training data. Models were trained using a Linux workstation equipped with 3 nvidia Quadro P6000 GPUs. Pix2pix models were trained for 200 epochs and vid2vid models were trained for 25 epochs. For validation, 35 image sequences from 3 held out specimens were reserved exclusively for performance metric calculations and were not used during training.

**SIGHT inference pipeline**

The workflow for synthetically immunolabeling entire OTLS datasets is outlined in **Supplementary Figure 2**. First, H&E analog channels are broken into 25 separate blocks (1024 x 1024 x 640 voxels in size with 25% lateral overlap). Each image block undergoes the same histogram adaptation for 16-bit to 8-bit conversion used to process the vid2vid training data, described above. 3D image volumes are separated into individual z levels for inference. Both *pix2pix* and *vid2vid* expect 3-channel input images for their data loading structures so the nuclear channel (TOPRO-3) is used as two input channels ch0 & ch1, eosin is used as the third ch2.

The first five levels of each block are processed using *pix2pix* to 'prime' the *vid2vid* network which synthetically labels the rest of the image block (**Supplementary Figure 2A**). Finally, the inferred image blocks are reassembled into a single 3D image volume using the stitching plugin from ImageJ. This procedure uses a snake-by-rows technique to stitch individual blocks together at the overlapping seams (via affine transformation) and then linear blending to remove any stitching artifacts (**Supplementary Figure 2B**).

**SIGHT heatmap generation**

The cancer heatmap algorithm uses the following logic sequence outlined in **Supplementary Figure 3**:

(A) The average signal level of the block for the hematoxylin analog channel is measured. If the signal falls below the background threshold for either channel, the block is ignored, and its score is set to zero. This step ensures that empty spaces, such as prostate gland lumens, are excluded.

(B) The number of pixels within the block containing CK8 signal above the background threshold is summed. If this count is less than 1% of the block's volume, the block is ignored, and its score is set to zero. This prevents the inclusion of blocks containing only stroma.

(C) For blocks that pass the above criteria, the score is calculated as:

$$P = 1 - \frac{\sum (CK5 > \beta_5)}{\sum (CK8 > \beta_8)}$$

where CK5 and CK8 represent the signal levels of each synthetic immunolabel within the block, and $\beta_5$ and $\beta_8$ denote their respective background thresholds.

The initial heatmap is coarse and the score of each block is independent of adjacent blocks (**Supplementary Figure 3B**). To smooth the heatmap a median filter with a cross-shaped structuring element is applied across each block. Next, scores between adjacent blocks are smoothed through linear interpolation followed by a gaussian filter ($\sigma$ = 25 voxels), ensuring smoother transitions across the dataset. The scoring procedure was implemented using the Python libraries *numpy* and *scikit-image*. For efficient processing, the framework leverages the automatic parallelization capabilities of the *dask* library. The heatmap generation step took approximately 20 minutes on a desktop computer equipped with 512 GB of RAM.

**3D Glandular segmentation**

Using a previously published deep-learning approach, trained a 3D segmentation to semantically label three key prostate microstructures—lumen, glandular epithelium, and stroma, directly from H&E analog images (**Supplementary Figure 4**). For model training, segmentation masks of each microstructure were generated by applying traditional computer vision techniques to OTLS data and the synthetically immunolabeled data created by vid2vid. For each training dataset: the glandular epithelium mask was obtained by applying Otsu's thresholding method to the synthetic CK8 channel. To acquire a solid epithelium mask with minimal holes/noise, first the synthetic CK8 image is blurred with a gaussian filter. A binary closing (dilation followed by erosion) is performed using a r=5 circular structural element, followed by a small-hole filling and small-object removal applied to the epithelium mask. Lumen masks were created using a percentile threshold for the $10^{th}$ percentile of the dynamic range of the eosin channel. Finally, the stroma segmentation mask was generated by combining the lumen and epithelium masks, subtracting those regions from the eosin channel and applying a triangle threshold. (**Supplementary Figure 4A**).

To automate segmentation, NN-UNet was implemented to train a model to predict these segmentation labels directly from the H&E analog channels of OTLS datasets (**Supplementary Figure 4B**). The model was trained for 1,000 epochs over 72 hours on a desktop computer equipped with 512 GB of RAM and a GeForce RTX 3090 GPU. On a held-out validation dataset, the final model achieved an average Dice coefficient of 0.89, demonstrating strong agreement with traditional computer vision-based segmentations. Following model training and validation the 3D gland segmentation model was used to segment prostate microstructures in OTLS datasets of intact 3 mm punch biopsies (**Supplementary Figure 4C**).

**Separation of benign and cancerous prostate glands**

SIGHT heatmaps are binarized to identify cancer-enriched regions within OTLS datasets through a thresholding approach. An optimal threshold value was determined by comparing the F1-score (Dice coefficient) of pathologist annotations of prostate glands and corresponding SIGHT heatmaps (Figure 4).  Binary masks of cancer-enriched regions are created by thresholding the SIGHT heatmap of each dataset at a threshold of 0.7. Finally, to separate the glands within the cancer-enriched regions of each dataset, the binarized masks are overlaid onto their corresponding 3D gland segmentation masks.

**Supplementary video descriptions**

**Supplementary Video 1. Nondestructive 3D pathology dataset of a 3 mm punch biopsy of prostate tissue imaged with OTLS**. OLTS enables imaging of diagnostically significant microstructures (such as glands and nuclei) with orders of magnitude greater sampling than standard 2D histology. Here the prostate biopsy is stained with rapidly diffusing small molecule fluorescent analogs of H&E and false colored to mimic the standard appearance of H&E histology.

**Supplementary Video 2. Synthetically immunolabelled 3 mm prostate biopsy**. Synthetic immunolabeling models trained using conditional-GANs can extract molecular information from H&E analog datasets to differentiate between different prostate microstructures. Here low molecular weight CK8 expressed in the luminal cell layer of prostate glands is shown in green, and high molecular weight CK5 expressed in the basal cell layer of benign prostate glands is shown in magenta. With TOPRO-3 (hematoxylin analog) shown in blue.

**Supplementary Video 3. 3D gland segmentations following application of SIGHT heatmap**. The glandular epithelium of a 3D pathology dataset with SIGHT heatmap coloring overlaid, where predicted cancerous glands are shown in red/orange hues, and benign glands are shown in blue/green.

**Supplementary Video 4. Depth stack visualization of SIGHT-generated heatmaps overlaid on OTLS microscopy data, false-colored to resemble H&E histology**. Warmer hues (red/orange) indicate regions predicted as cancerous, while cooler tones (blue/green) denote benign tissue. SIGHT enables rapid and accurate detection of cancerous prostate microstructures across large 3D volumes.

**Supplementary Video 5. Classification of cancerous and benign prostate glands**. Combining the 3D gland segmentations with the binary segmentation mask of the cancer-enriched regions predicted by SIGHT allow us to classify prostate microstructures as either cancerous (red) or benign (blue). Separation of benign and cancerous structures at a glandular level is fully automated more precise than our previous 3D pathology studies. This automation enables large-scale quantitative analysis and classification of clinically relevant microstructures and may facilitate the discovery of novel prognostic biomarkers in prostate cancer.

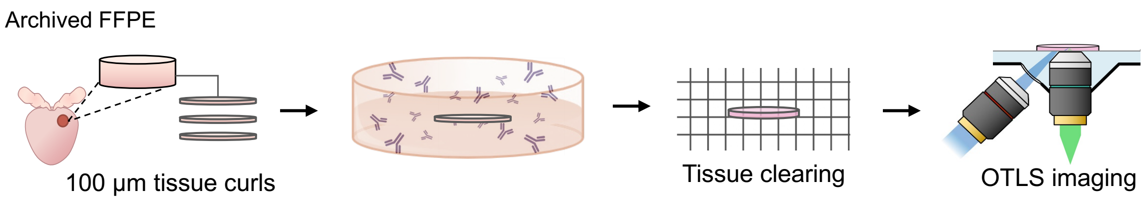


| Time | Main purpose | Detailed steps |
|---|---|---|
| **Week 1** | | RT = Room Temperature, O/N = overnight |
| Day 1 | Dehydration | Wash with PBS-to-methanol series at room temperature (RT). |
| Day 2 | Bleaching | Chill at 4°C, then bleach in 5% $H_2O_2$ in methanol at 4°C overnight (O/N). |
| Day 3 | Rehydration **+ cytoplasm stain** | Wash in methanol-to-PBS series at RT. Incubated in Alexa Fluor™ 488 NHS ester (in pH 5 PBS) at 37°C O/N. |
| Day 4 | Washing | Wash in PBS/Triton X-100 for 1h at RT. |
| **Week 2** | | |
| Day 1 | Reduce background | Incubate in PBS/Triton X-100/DMSO/Glycine at 37°C O/N. |
| Day 2 | Blocking | Block with PBS/TritonX-100/DMSO/Donkey Serum + $NaN_3$ at 37°C O/N. |
| Day 3 | Primary antibody staining | Incubate with the primary antibody (CK8 or CK5 Monoclonal Antibody) in PBS-Tween with Heparin/DMSO/Donkey Serum + $NaN_3$ at 37°C for 2 days. |
| Day 5 | Washing | Wash in PBS-Tween with Heparin at RT. |
| Day 6 | Secondary antibody staining | Incubate with the secondary antibody (Alexa Fluor 647 Donkey Anti-Mouse IgG) in PBS-Tween with Heparin/Donkey Serum + $NaN_3$ at 37°C for 2 days. |
| **Week 3** | | |
| Day 1 | Washing **+ nuclei stain** | Wash in PBS-Tween with Heparin at RT. Incubate with SYTO™ 85 O/N at 37°C. |
| Day 2 | Dehydration | Wash with PBS, then incubate with DI-water-to-ethanol series at RT. |
| Day 3 | **ECi clearing** | Incubate in Ethyl Cinnamate (ECi) at RT. Ready for 3D imaging. |

**Supplementary Figure 1. Tri-labeling protocol for OTLS datasets.**

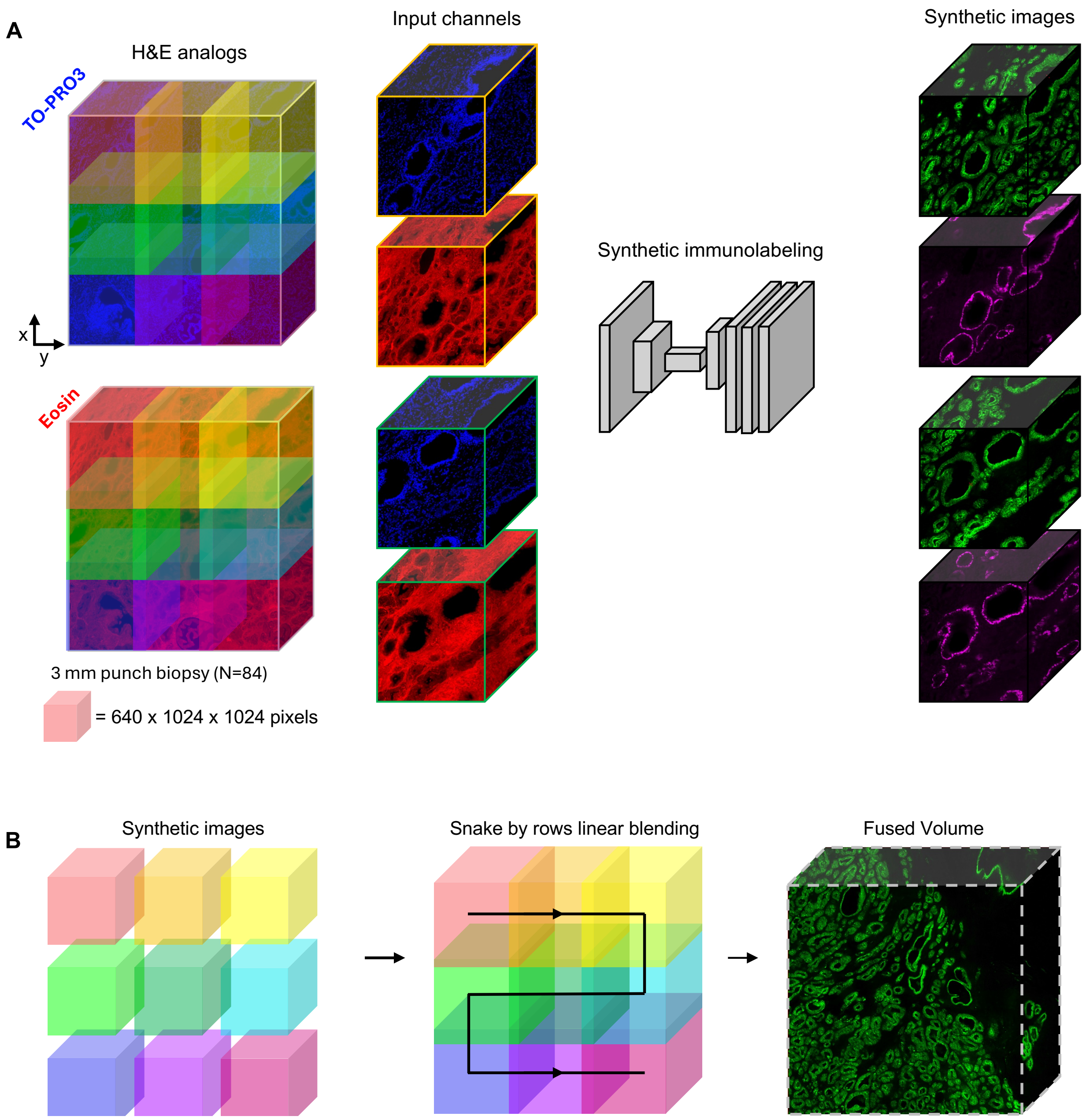


**Supplementary Figure 2. Synthetic immunolabeling inference workflow. (A)** Preprocessing. H&E analog OTLS images of 3 mm PCa biopsies are broken into separate 1024 x 1024 z-stacks with 25% overlap between adjacent blocks. Individual levels are saved as separate image files and saved to disk in a format compatible with the vid2vid data loader. **(B)** Post processing. Synthetic image sequences are generated from vid2vid as separate 1024 x 1024 sized z-stacks. Adjacent image sequences are stitched together using an affine transformation and linear blending to remove artifacts generated at the seams of adjacent blocks, creating a final synthetically labelled volume.

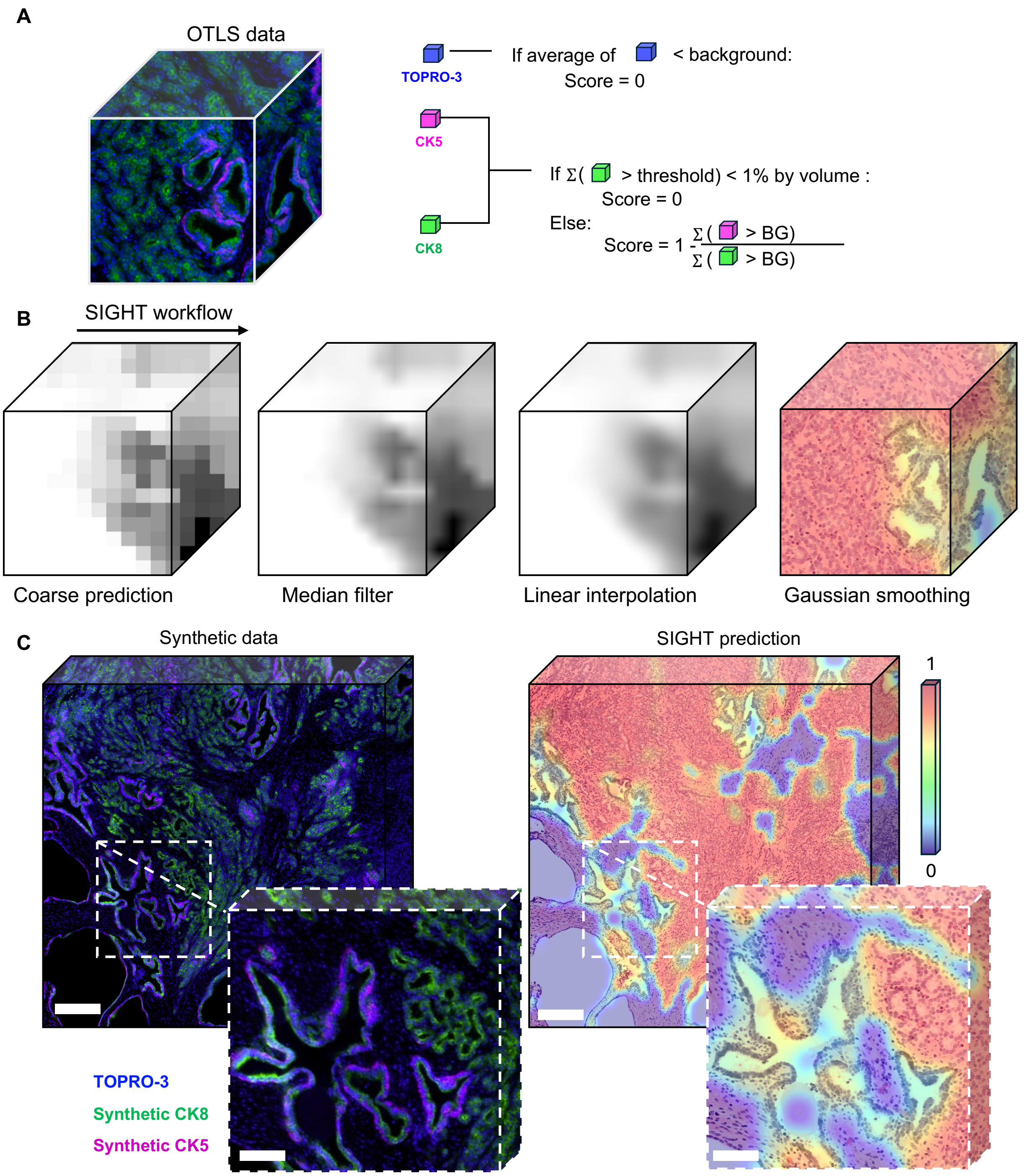


**Supplementary Figure 3. Heatmap generation from synthetic immunolabels. (A)** Following synthetic immunolabeling, OTLS datasets are subdivided into blocks measuring 50 µm a side. Within each block the ratio of CK5 to CK8 signal above the background threshold is measured and subtracted from 1. **(B)** The resulting heatmap is then smoothed using a series of filters. **(C)** Using this process a heatmap of the cancer enriched regions of a 3D pathology dataset of a 3 mm punch biopsy can be generated in 25 minutes. SB = 300 & 100 µm

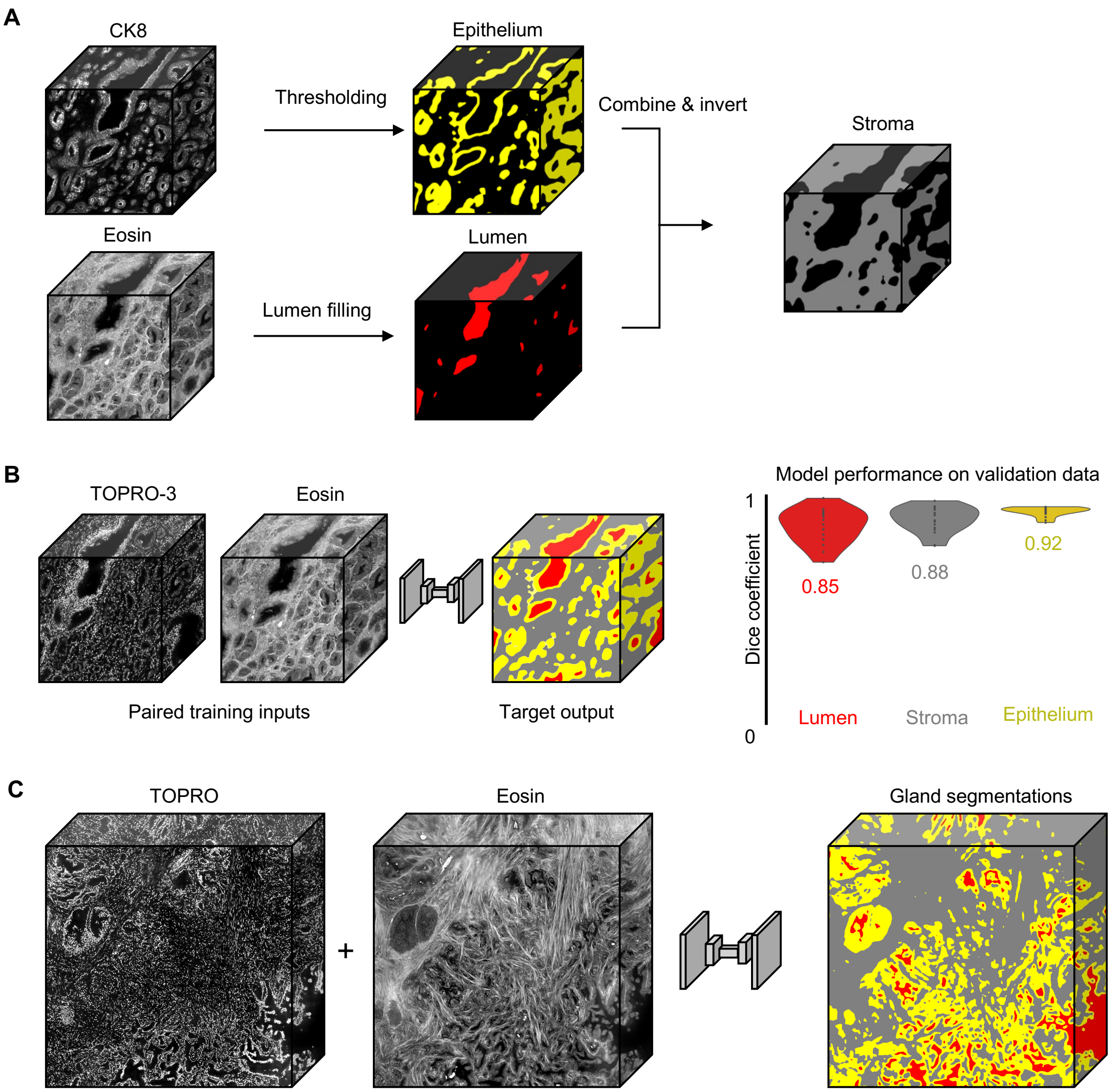


**Supplementary Figure 4. 3D Gland Segmentation Workflow. (A)** Traditional computer vision techniques are applied to synthetic CK8 and eosin channels to generate training labels. The glandular epithelium was segmented by thresholding the CK8 channel, and lumen regions were identified by filling holes in the eosin channel. These masks were then combined and subtracted from the eosin channel, and a threshold was applied to obtain the stroma mask. **(B)** The training labels from Panel A were used to train a 3D semantic segmentation model using nnU-Net. The model was trained to predict three prostate microstructures—lumen, epithelium, and stroma—directly from H&E analog channels. Performance on held-out validation data demonstrated high accuracy based on the Dice coefficient. **(C)** The trained segmentation model was applied to a 3 mm prostate punch biopsy to generate a full-resolution 3D semantic segmentation of the three key microstructural compartments: lumen, epithelium, and stroma.

| Patient ID | BCR category | Racial | Time to BCR (Days) | ISUP Group |
|---|---|---|---|---|
| afm001 | 0 | AF | 90 | 2 |
| afm004 | 0 | AF | 4253 | 1 |
| afm005 | 0 | AF | 754 | 1 |
| afm008 | 0 | AF | 4451 | 2 |
| afm009 | 0 | AF | 193 | 1 |
| afm010 | 0 | AF | 299 | 2 |
| afm012 | 1 | AF | 2069 | 1 |
| afm014 | 1 | AF | 3606 | 1 |
| afm016 | 0 | AF | 377 | 2 |
| afm021 | 0 | AF | 3061 | 1 |
| afm023 | 0 | AF | 3074 | 2 |
| afm024 | 0 | AF | 4426 | 1 |
| afm027 | 0 | AF | 363 | 2 |
| afm033 | 0 | AF | 5704 | 1 |
| afm035 | 1 | AF | 666 | 5 |
| afm039 | 1 | AF | 278 | 1 |
| afm042 | 1 | AF | 1246 | 1 |
| afm047 | 1 | AF | 2261 | 2 |
| afm050 | 1 | AF | 2038 | 5 |
| afm052 | 1 | AF | 1070 | 2 |
| afm054 | 0 | AF | 2857 | 1 |
| afm059 | 0 | AF | 2920 | 1 |
| afm066 | 1 | AF | 590 | 1 |
| afm069 | 0 | AF | 1923 | 1 |
| afm072 | 0 | AF | 102 | 2 |
| afm073 | 1 | AF | 1316 | 3 |
| afm074 | 1 | AF | 152 | 2 |
| afm076 | 1 | AF | 374 | 2 |
| afm077 | 0 | AF | 461 | 3 |
| afm078 | 1 | AF | 32 | 5 |
| afm080 | 0 | AF | 1782 | 2 |

| | | | | |
|---|---|---|---|---|
| afm081 | 0 | AF | 1870 | 3 |
| afm082 | 0 | AF | 1840 | 2 |
| afm086 | 1 | AF | 783 | 2 |
| afm090 | 0 | AF | 3539 | 3 |
| afm096 | 0 | AF | 719 | 1 |
| afm097 | 0 | AF | 5096 | 2 |
| afm098 | 0 | AF | 603 | 2 |
| afm100 | 0 | AF | 648 | 1 |
| afm101 | 0 | AF | 3148 | 1 |
| afm189 | 0 | AF | 1590 | 2 |
| eam002 | 1 | CA | 699 | 3 |
| eam003 | 0 | CA | 4144 | 2 |
| eam006 | 1 | CA | 307 | 3 |
| eam009 | 1 | CA | 123 | 2 |
| eam017 | 0 | CA | 3356 | 2 |
| eam018 | 1 | CA | 838 | 1 |
| eam020 | 1 | CA | 791 | 3 |
| eam022 | 0 | CA | 91 | 1 |
| eam025 | 0 | CA | 477 | 2 |
| eam033 | 1 | CA | 930 | 1 |
| eam034 | 1 | CA | 339 | 4 |
| eam035 | 0 | CA | 606 | 2 |
| eam037 | 0 | CA | 1044 | 1 |
| eam041 | 1 | CA | 665 | 2 |
| eam042 | 0 | CA | 298 | 2 |
| eam043 | 1 | CA | 73 | 4 |
| eam044 | 0 | CA | 2458 | 1 |
| eam047 | 0 | CA | 1200 | 1 |
| eam049 | 0 | CA | 462 | 1 |
| eam056 | 0 | CA | 1125 | 1 |
| eam057 | 0 | CA | 2501 | 1 |
| eam060 | 1 | CA | 529 | 2 |

| | | | | |
|---|---|---|---|---|
| eam062 | 1 | CA | 268 | 1 |
| eam069 | 0 | CA | 3495 | 3 |
| eam071 | 0 | CA | 541 | 1 |
| eam076 | 0 | CA | 365 | 2 |
| eam078 | 0 | CA | 1993 | 2 |
| eam083 | 0 | CA | 2003 | 2 |
| eam085 | 0 | CA | 952 | 5 |
| eam088 | 1 | CA | 1050 | 2 |
| eam089 | 0 | CA | 85 | 2 |
| eam092 | 0 | CA | 95 | 1 |
| eam098 | 1 | CA | 1496 | 2 |
| eam100 | 1 | CA | 1302 | 4 |

**Supplementary Table 1. Clinical parameters for patient cohort.**

BCR category: 1 = cases with BCR under 5 years, 0 = all other cases

| **Feature Category** (G: gland, E: epithelium, L: lumen, S: stroma) | **3D Glandular Features** |
|---|---|
| **Size** | Volume (L) / volume (E)<br><br>Volume (E) / volume (G)<br><br>Volume (S) / volume (G) |
| **Compactness** | Surface area (G) / volume (G)<br><br>Surface area (E) / volume (E)<br><br>Surface area (L) / volume (L) |
| **Irregularity** | Volume (G) / convex hull volume (G)<br><br>Volume (E) / convex hull volume (E)<br><br>Volume (L) / convex hull volume (L) |
| **Boundary Curvature** | Average Gaussian curvature (absolute values) of the G surface<br><br>Average Gaussian curvature (absolute values) of the E surface<br><br>Average Gaussian curvature (absolute values) of the L surface |

**Supplementary Table 2. List of glandular morphological features used for prognostic analysis**
